# Protein Structure Prediction Using a Maximum Likelihood Formulation of a Recurrent Geometric Network

**DOI:** 10.1101/2021.09.03.458873

**Authors:** Guowei Qi, Mallory R. Tollefson, Rose A. Gogal, Richard J. H. Smith, Mohammed AlQuraishi, Michael J. Schnieders

**Author notes:** The authors contributed equally (Guowei Qi, Mallory R. Tollefson).

## Abstract

Only ∼40% of the human proteome has structural coordinates available from experiment (*i*.*e*., X-ray crystallography, NMR spectroscopy, or cryo-EM) or homology modeling with quality templates (*i*.*e*., 30% sequence identity or greater), leaving most of the proteome structurally unsolved. Deep learning (DL) methods for predicting protein structure can help close knowledge gaps where experimental and homology models are difficult to obtain. Recent advances in these DL methods have shown promising results in expanding structural coverage to the scale of the entire human proteome, providing researchers with more complete protein structural information. Here, we improve upon an existing DL algorithm for protein structure prediction, the Recurrent Geometric Network (RGN). We first expand the training dataset to include experimental uncertainty data in the form of atomic displacement parameters, then derive a maximum likelihood loss function that incorporates this uncertainty data into model training. Compared to the original RGN, our novel maximum likelihood model improves the rate of convergence of initial model training and ultimately results in more accurate structure prediction according to the root mean square deviation (RMSD) of backbone atoms, the Global Distance Test (GDT), the Global Distance Test High Accuracy (GDT-HA), and the Template-Modeling Score (TM-Score). Our model also predicts structures with more favorable backbone torsions, which provide more accurate starting coordinates for downstream physics-based simulations. Based on these results, our maximum likelihood reformulation provides a framework for improving existing or future machine learning algorithms for protein structure prediction. The augmented dataset, data collection scripts, reformulated RGN source code, and a series of trained models are publicly available at https://github.com/SchniedersLab/likelihood-rgn.

## Introduction

Nearly 180,000 biomolecular structures obtained using experimental techniques, such as X-ray crystallography, nuclear magnetic resonance (NMR) spectroscopy, or cryogenic electron microscopy (cryo-EM), are available within the Protein Data Bank (PDB)(1), yet the majority of the human proteome lacks structural coverage. Only ∼40% of the human proteome has been structurally solved through either experimental methods or homology modeling using templates with greater than 30% sequence identity(2). The fold of a protein can be used to assist in understanding protein function and identifying potential drug therapy targets. For these reasons, the lack of structural coverage for the human proteome is a central problem in biochemistry(3). Computational methods can supplement the structural data available from experiment, and recent advances in such methods(4-6) have increased the feasibility of predicting a protein’s structure from its primary amino acid sequence (*i*.*e*., the protein folding problem)(7).

Traditional computational methods for predicting a protein fold combine a physics-based model of intermolecular forces(8-16), an explicit(17) or continuum(8-16, 18-20) solvation model, and a sampling algorithm(5, 21) that builds upon molecular dynamics simulations. However, a limiting factor of a physics-based approaches is that protein folding often occurs on millisecond (10^−3^ seconds) or longer timescales, whereas GPU-accelerated molecular dynamics simulations are largely limited to microsecond (10^−6^ seconds) time scales due to the computational expense of computing interactions over all atoms in a protein system.

Advances in machine learning—specifically, deep learning (DL)—have prompted the development of new data-driven approaches to predicting protein structure(22, 23). These DL algorithms use existing protein data, such as 3D coordinates, evolutionary data, and multiple sequence-alignments (MSAs), to train a computational model that predicts protein structure from an amino acid sequence. Two benefits these DL methods provide are 1) faster predictions after model training compared to physics-based protein folding methods and 2) predicted structures that can be used for downstream computational analyses (*e*.*g*., free energy perturbation, docking, *etc*.) in cases where coordinates from experiment or homology remain unavailable. These data-driven solutions to the protein folding problem have become so significant that comprehensive, homogenous datasets of protein structures have been curated specifically for machine learning(24, 25).

The success of DL methods for protein structure prediction was demonstrated by the DeepMind(26) research group at the 14^th^ Critical Assessment of Structure Prediction (CASP14) competition using their AlphaFold2 algorithm(27). At CASP14, AlphaFold2 achieved a median Global Distance Test (GDT) of 92.4, corresponding to an average root-mean-square-deviation (RMSD) error of 1.6 Å. AlphaFold2 was also recently used to predict structures for 98.5% of the human proteome, attaining confident predictions in 58% of all residues(28). Another DL model, the Recurrent Geometric Network (RGN)(29), uses end-to-end differentiable learning of protein structure and was able to predict coordinates within 1.5 Å RMSD of the top performing servers at the 12^th^ CASP competition (CASP12) of 2016. RGN predicts three torsions (*i*.*e*., ϕ, ψ, and ω) for each amino acid in a protein sequence and sequentially builds the complete backbone by computing internal coordinates from known bond lengths and bond angles (Figure 1). Most recently, the second iteration of RGN—termed RGN2—was shown to successfully predict protein structure from single sequences without the use of MSAs(30). RGN2 outperformed both AlphaFold2 and RoseTTAFold(31) in predicting the structures of orphan proteins while also achieving a 10^6^-fold reduction in compute time.

**Figure 1.**
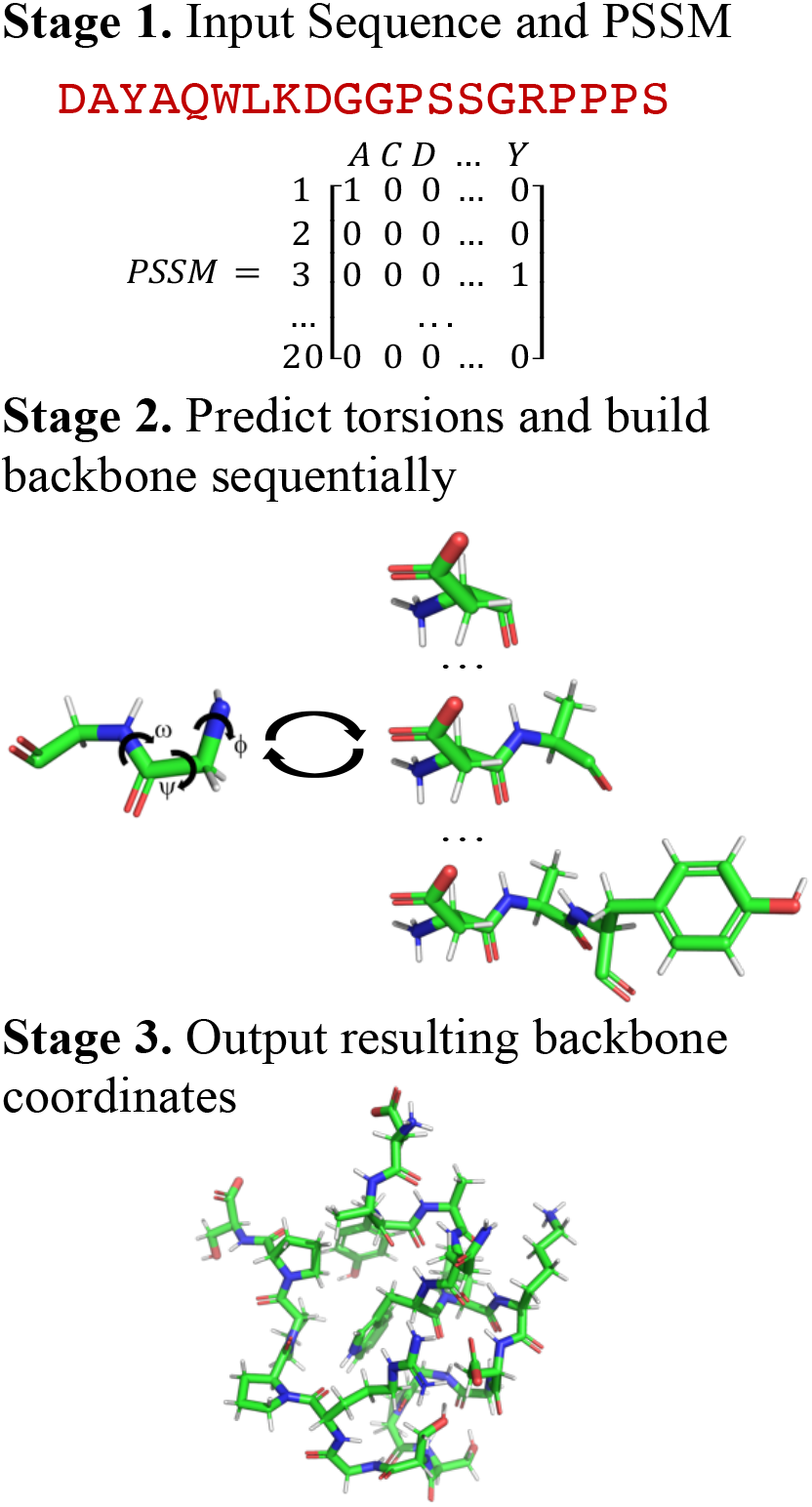
The three stages of RGN. In stage 1, a primary sequence and Position Specific Scoring Matrix (PSSM) are submitted to the RGN; in stage 2, three backbone torsions (*i*.*e*., ψ, ω, and ϕ) are predicted for each amino acid and the backbone is sequentially built; stage 3 outputs the final 3D structure and computes the loss (*i*.*e*., dRMSD between the predicted structure and experiment).

Further improvements to these DL approaches will help facilitate structure prediction for new amino acid sequences, nucleic acids, and their complexes. Currently, most DL models for structure prediction, including RGN, RGN2, and AlphaFold2 are trained and evaluated using a least-squares style target function (*e*.*g*., RMSD between the predicted structure and the experimental structure). This target function trains the neural network as if all atomic coordinates within a structure are equally certain, which is not the case for experimentally determined structures due to factors such as the intrinsic flexibility of the protein and the overall quality of the experiment.

In fields such as experimental biology, structural refinement is performed using a maximum likelihood approach, where the target function is modified to account for the uncertainty of each atomic coordinate prior to optimization. The use of these maximum likelihood approaches in experimental structural biology has a rich history(32, 33), which includes application to X-ray crystallography refinement(34, 35), molecular replacement(36-38), and NMR refinement(39, 40). For example, the initial application of a maximum likelihood target function to X-ray refinement in *XPLOR(41)* achieved more than twice the improvement to average phase error compared to least-squares refinement(34). In the context of molecular replacement in the program *Phaser*(38), likelihood-enhanced rotation and translation targets were shown to be more sensitive to the correct orientation and translation, respectively, than the corresponding Crowther fast rotation function(36) and correlation-coefficient fast translation function(37). Finally, using maximum likelihood and Bayesian principles for NMR refinement resulted in structures that were optimal in terms of accuracy and structural quality(40).

Here, we derive and implement a new DL model, Likelihood-RGN, which applies the principle of maximum likelihood refinement to the original RGN model to improve protein structure prediction. Structures available in the PDB vary in quality (the majority were determined based on a resolution worse than 2 Å(42)) and often exhibit different degrees of disorder among the regions of even a single protein domain. Using a maximum likelihood target to train a neural network allows higher quality regions of structures with more certainty in atomic coordinates to have a greater impact on model training, while poorer quality or more disordered regions of structures contribute to a lesser extent. To accomplish this, we first compile an improved, homogenous training dataset, termed the ProteinNetX, which includes B-factors from X-ray crystallography and computed atomic displacement parameters from NMR spectroscopy as measures of experimental uncertainty. Following the generation of the dataset, we derive a maximum likelihood loss function based on the electron density function from X-ray crystallography that incorporates uncertainty data into model training. We train a new model, Likelihood-RGN, using this maximum likelihood loss function and generate predictions for a series of target structures from CASP12. We compare our predicted structures to experiment, showing significant improvement over structures predicted by the original (least-squares loss) RGN according to several global distance and geometry metrics, such as the RMSD of backbone atoms, the Global Distance Test (GDT)(43), the Global Distance Test High Accuracy (GDT-HA)(44), the Template-Modeling Score (TM-Score)(45), and the proportion of favored and outlying torsions. Finally, we perform physics-based optimizations on structures predicted by both RGN and Likelihood-RGN to determine their respective suitability for downstream computational analyses. These improved results strongly suggest that a maximum likelihood approach can be incorporated into more advanced DL models, such as RGN2 and AlphaFold2, in order to generate increasingly accurate predictions of protein structure.

## Results

### A. The ProteinNetX Structure Prediction Dataset with Temperature Factors

When limited to protein structures solved by X-ray crystallography with experimental B-factors, ProteinNetX contains 84.7% of the structures included in the original CASP12 ProteinNet after filtering the dataset at 90% sequence identity. After including multi-model NMR structures with computed atomic displacement parameters, ProteinNetX covers 94.8% of the structures available in the original CASP12 ProteinNet dataset (Figure 2a). When computing NMR atomic displacement parameters, we add a constant of 20 Å^2^ to each value so that the peak of the distribution (Figure 2c) shifts to match the peak of the distribution of crystallographic B-factors (Figure 2b). This shift results in a final distribution of atomic displacement parameters (Figure 2d) that mirrors the distribution of experimental temperature factors. Computing atomic displacement parameters for single-model NMR and cryo-EM structures is beyond the scope of this work but may be explored in the future.

**Figure 2.**
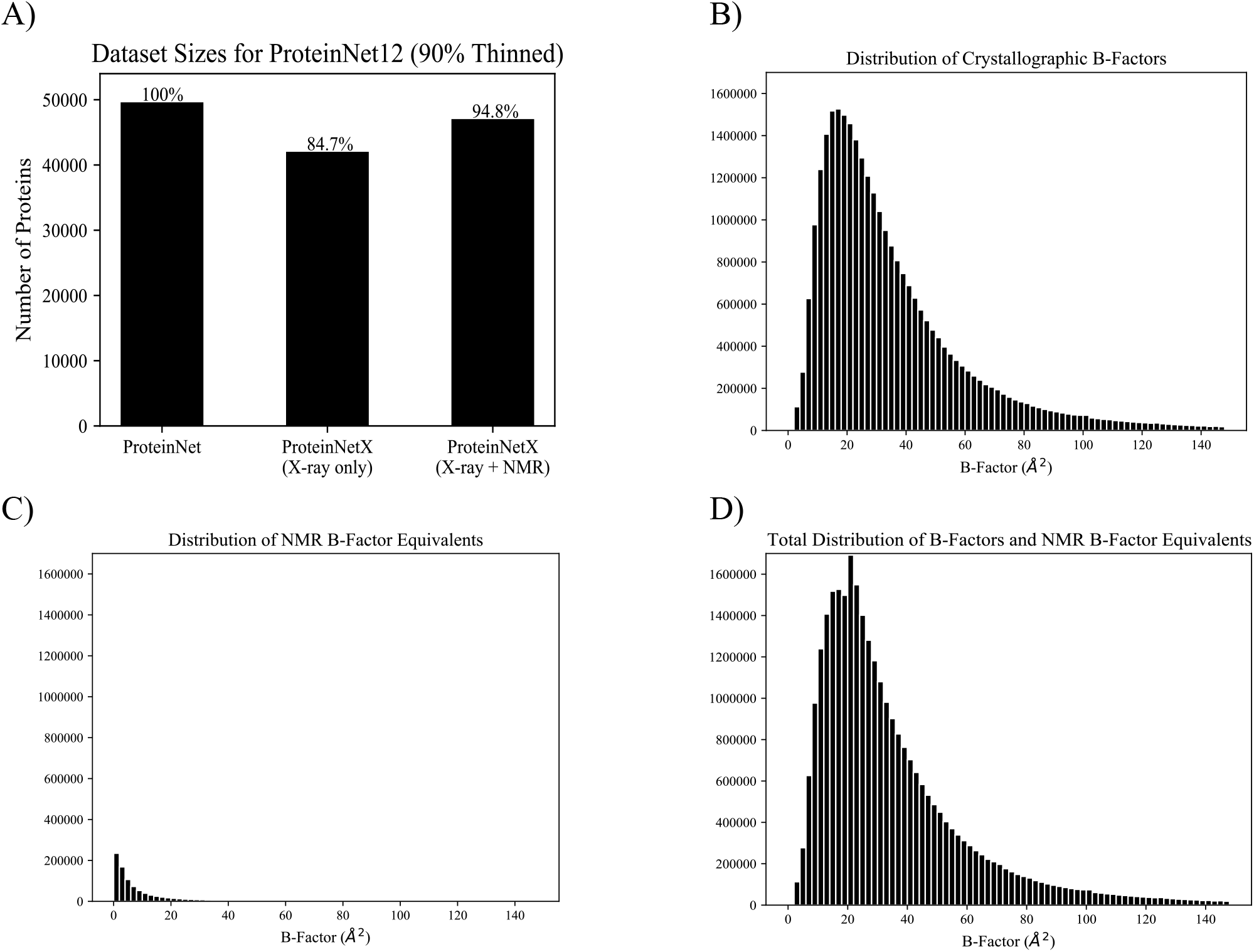
**A)** Dataset sizes for three variations of ProteinNet. Left shows the ProteinNet dataset as originally published (49,600 structures), middle shows the ProteinNetX dataset with only structures from X-ray crystallography (42,019 structures), and right shows the full ProteinNetX dataset with multi-model NMR structures included (47,035 structures). **B)** Distribution of crystallographic B-factors for all X-ray structures in ProteinNetX. **C)** Distribution of computed atomic displacement parameters for multi-model NMR structures in ProteinNetX. **D)** Distribution of all atomic displacement parameters in the full ProteinNetX dataset.

### B. Improved Structure Prediction with a Maximum Likelihood Loss

We first trained five pairs of models using the CASP12 ProteinNetX dataset containing only X-ray crystallography structures. Each pair was initialized from the same originally published hyperparameters(29) and random seed while controlling for all factors aside from the loss function. Within each pair, one model was trained using the original, least-squares loss function (*i*.*e*., dRMSD) and the other model was trained using our maximum likelihood loss. Each model was trained for 1.5 million iterations (where one iteration of training occurs over a batch of 32 proteins), which was followed by an additional 10,000 iterations of training at a reduced learning rate to achieve a small but noticeable gain in prediction accuracy. Plotting a running mean of the average dRMSD of structures in the testing dataset over the training iterations for each trial reveals that using a maximum likelihood loss to reduce the contributions of highly disordered regions of proteins results in smoother convergence during initial training (Figure 3a). Similarly, using a maximum likelihood loss results in an improved final convergence compared to using the least-squares, dRMSD loss (Figure 3c).

**Figure 3.**
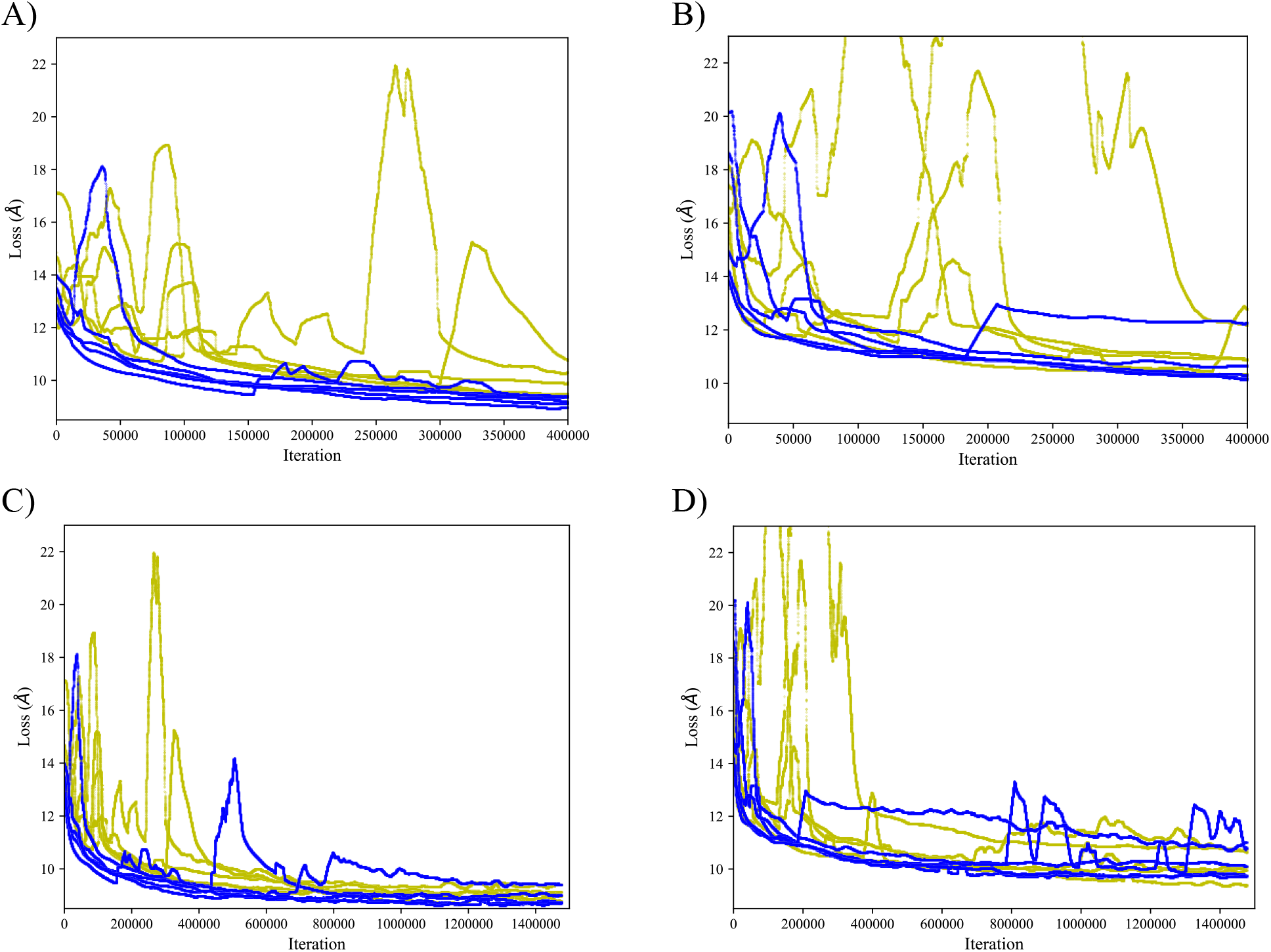
Running average least-squares loss of the testing dataset. Five trials of the first 400,000 training iterations using ProteinNetX with **A)** only X-ray structures and **B)** X-ray and NMR structures show that training with a maximum likelihood loss (blue curves) results in smoother gradient descent over the initial training period compared to training with the least-squares loss (yellow curves), as the network places less weight on highly disordered regions of proteins when B-factors are included in the loss function. Training over 1,500,000 iterations with **C)** only X-ray structures and **D)** both X-ray and NMR structures shows that the maximum likelihood RGN models (blue curves) tend to converge to a smaller value than the least-squares RGN models (yellow curves).

This same procedure was used to train models using the full ProteinNetX dataset (*i*.*e*., including x-ray and multi-model NMR structures) generated from the 90% thinned CASP12 ProteinNet. Using the full ProteinNetX dataset, the maximum likelihood model again showed increased stability during initial training (Figure 3b) and improved convergence (Figure 3d) when compared to the original, least-squares model trained on the same set of proteins.

Using the final trained models resulting from each trial, predicted structures were generated for a testing dataset of 63 CASP12 target structures not present in the training dataset(24). When comparing the models that generated the most physically realistic protein structures (*i*.*e*., those with the largest proportion of favored backbone torsions) from each set of trials, Likelihood-RGN outperforms the original RGN when evaluating the global accuracy of the testing set structures using dRMSD, RMSD, GDT(43), GDT-HA(44), and TM-Score(45) (Table 1). This suggests our maximum likelihood model achieves improved global folding accuracy while maintaining local secondary structure. This increased predictive accuracy remains evident when we average over all five trials (Table S1), as well as when we train and evaluate models using the CASP11 training and testing sets (Table S3).

**Table 1.** Average scores for 63 testing set proteins across the best performing trials for the RGN (least-squares loss) and Likelihood-RGN (maximum likelihood loss) DL models. The neural networks here were trained on both the X-ray only ProteinNetX dataset and the full ProteinNetX dataset consisting of X-ray and NMR protein structures.

| Training Dataset | Model | dRMSD | RMSD | GDT | GDT -HA | TM-Score | Outlier Torsions | Favored Torsions |
| --- | --- | --- | --- | --- | --- | --- | --- | --- |
| X-Ray | RGN | 8.87 | 14.14 | 0.19 | 0.09 | 0.32 | 52.7 | 23.9 |
|  | L-RGN | 8.61 | 13.66 | 0.21 | 0.10 | 0.33 | 42.6 | 40.8 |
| X-Ray+NMR | RGN | 9.24 | 16.53 | 0.14 | 0.06 | 0.24 | 23.3 | 48.1 |
|  | L-RGN | 8.81 | 14.68 | 0.18 | 0.09 | 0.30 | 6.6 | 77.8 |

When examining the two models trained by the full ProteinNetX dataset, the original RGN converged to a minimum that predicted global protein topology with reasonable accuracy, but failed to predict local secondary structure. When using our maximum likelihood loss, the final trained Likelihood-RGN model converged to a minimum that predicted structures with a higher proportion of favored backbone torsions (Table 1). This increase in favored backbone torsions shown by the trained Likelihood-RGN models leads to a substantial improvement in local secondary structure prediction over the original RGN. Structures for six selected CASP12 targets demonstrate this improvement, both prior to physics-based optimization (Figure 4) and after physics-based optimization (Figure S1). The six selected targets show a variety of structures with different lengths (ranging from 89 to 409 amino acids) and varying secondary structure characteristics (*i*.*e*., both alpha helices and beta sheets). These targets demonstrate that Likelihood-RGN generally predicts alpha helices more accurately than beta sheets, while the original RGN fails to reproduce either form of secondary structure. The physically realistic structures output by Likelihood-RGN are more amenable to downstream physics-based optimization and thus, are more useful for further computational analyses.

**Figure 4.**
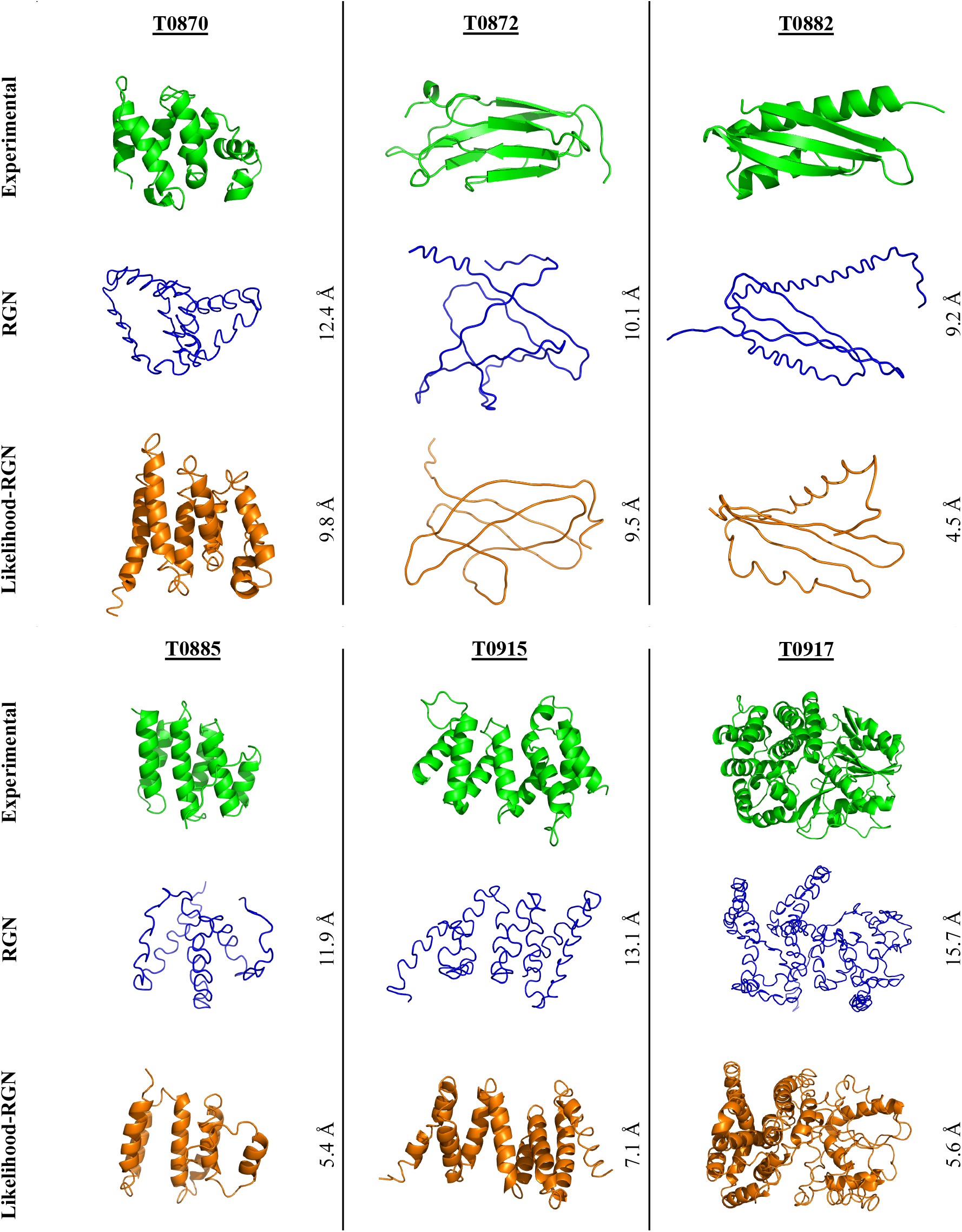
Six targets from the CASP12 competition shown in their experimentally solved coordinates (green), the original RGN predicted coordinates (blue), and the Likelihood-RGN predicted coordinates (orange) from the training trial that provided the best backbone torsions and corresponding RMSDs to the experimentally known structure. These structures are output directly from the machine learning model and have undergone no additional physics-based optimization. The Likelihood-RGN structures have a smaller RMSD to the known experimental fold compared to the original RGN.

Though adding NMR structures to the CASP12 training dataset improved the backbone torsions predicted by the final model, the global distance metrics were slightly worse compared to the initial set of models trained using only X-ray crystallography structures. While X-ray diffraction data represent both short- and long-range interatomic distances, NMR experimental observations mainly capture local interactions, which may explain the improvement in secondary structure prediction but worsening of the global distance metrics. Representation of NMR models in the dataset could be modified to improve upon global distance metrics in addition to predicted backbone torsions. For example, the entire NMR ensemble could be used in model training, rather than only providing the coordinates of the first structure of the ensemble. Future work could include investigating alternative methods to represent NMR coordinates and their uncertainty.

### C. Physics-Based Optimization of Predicted Backbone Structures

To determine the suitability of our predicted structures for downstream computational analyses, we performed a series of physics-based optimizations on the structures generated by both the RGN and Likelihood-RGN models trained using the full ProteinNetX dataset. The 63 testing set proteins were minimized using the Amber(12, 13) fixed charge and AMOEBA(15, 46) polarizable force fields with a generalized Kirkwood implicit solvent. Our minimization protocol causes a small increase in the average RMSD of the predicted structures. While the average favored and outlier torsions worsen for Likelihood-RGN during minimization (Table 2), Ramachandran plots show that the torsional angles for many test proteins disperse across favored regions more realistically compared to the clustered angles that are output directly from the DL network (Figure 5). After minimization, the structures predicted by Likelihood-RGN continue to show an improved RMSD, GDT, GDT-HA and TM-Score compared to the original RGN (Table 2) and the proportion of favored and outlier torsions in Likelihood-RGN continues to significantly surpass RGN, suggesting that the Likelihood-RGN structures will better retain their folds upon downstream biophysical analyses. Ramachandran plots (Figure 5) for six proteins from the testing set show that Likelihood-RGN consistently has fewer torsion outliers and more favored torsions than RGN, both before and after minimization.

**Table 2.**
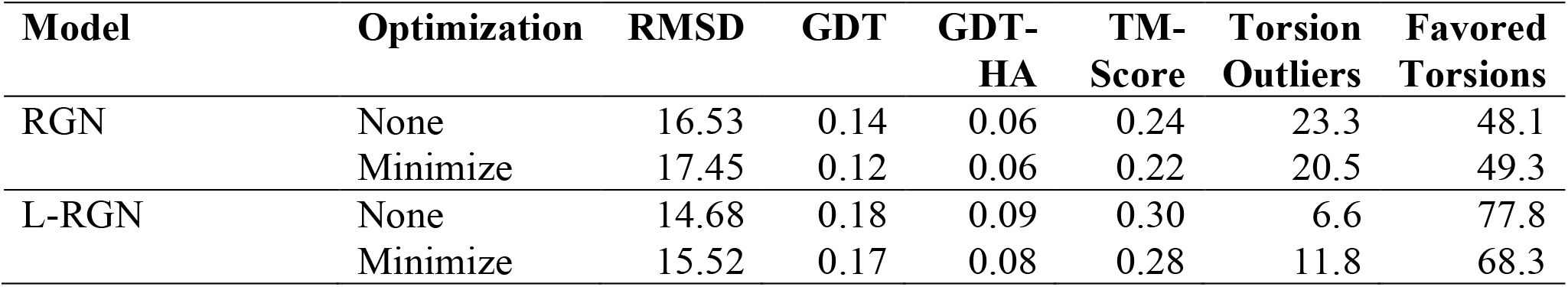
Average scores over the best trials for the 63 testing set protein structures directly predicted by both RGN and Likelihood-RGN, as well scores for the structures following minimization under the AMOEBA force field. This data was collected from trials that were trained using the X-ray+NMR ProteinNetX dataset.

**Figure 5.**
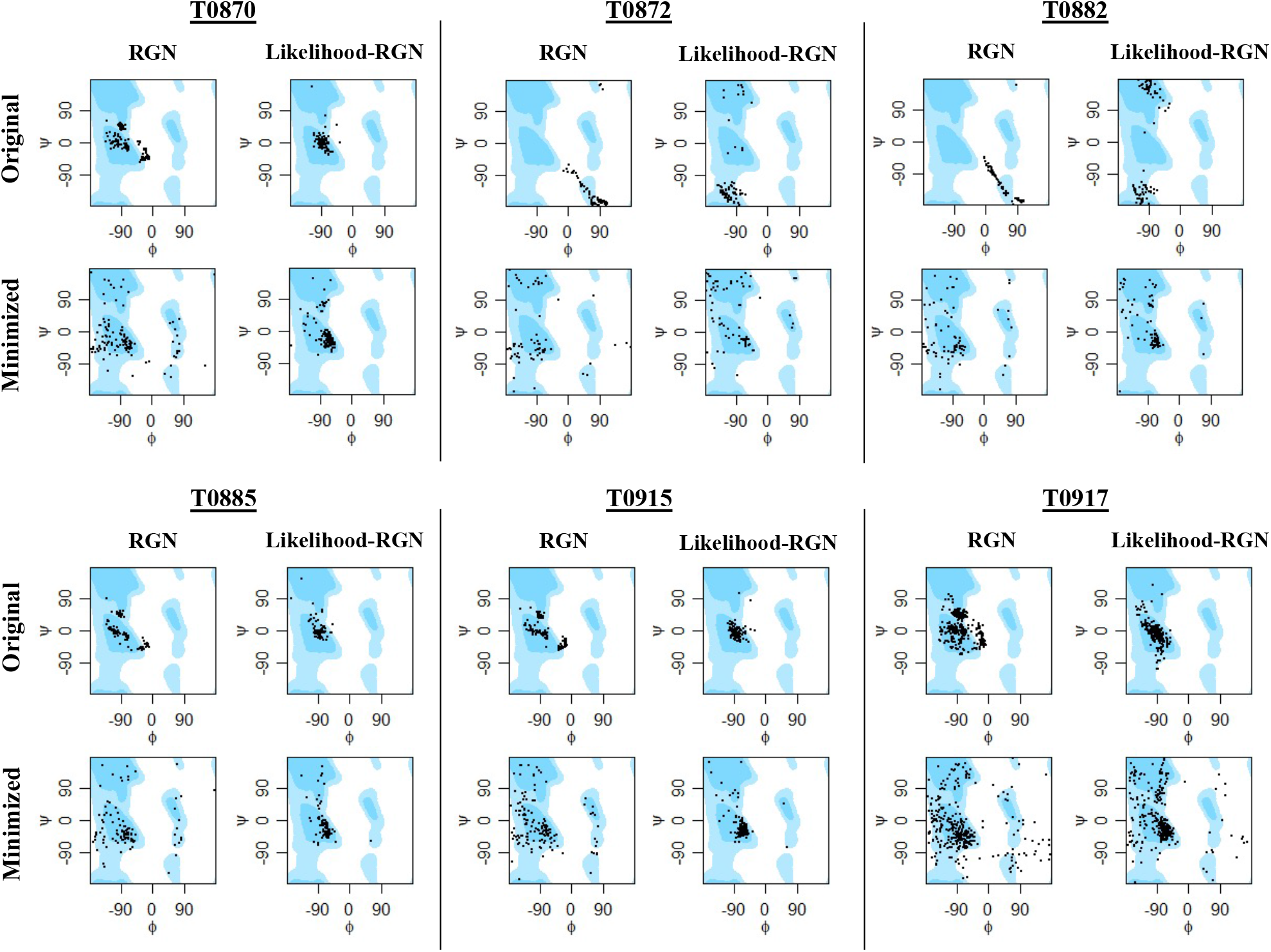
Ramachandran plots for the structures of six CASP12 targets as predicted by RGN and Likelihood-RGN. Ramachandran plots are also shown for the six targets after minimization with the AMOEBA force field.

After minimizing the 63 testing set protein structures, we applied our many-body sidechain repacking algorithm(47) to each structure. The RGN outputs only the backbone coordinates for each protein; therefore, applying our algorithm finalizes each structure by building sidechain atoms and placing the sidechains in their global minimum energy conformation. After optimizing sidechain placement in each of the testing set structures, we assessed quality of the structures prior to and after sidechain optimization using the heuristic MolProbity(48, 49) algorithm. MolProbity is used widely by crystallographers to aid refinement of x-ray structures and to evaluate the structures for steric clashes, rotamer placement, and favorable backbone torsions. The MolProbity algorithm provides a score that is calibrated to predict the resolution for an x-ray structure (*i*.*e*., a MolProbity score of 1.0 corresponds to an x-ray resolution of 1.0 Angstroms). Lower MolProbity scores are consistent with higher quality x-ray diffraction data. Optimizing sidechain placement of the output structures from RGN and Likelihood-RGN substantially improves the MolProbity score, clash score (*i*.*e*., the number of steric clashes per 1000 atoms), and favored torsions, and reduces the percentage of torsion and rotamer outliers (Table 3).

**Table 3.** Refinement statistics for the 63 testing set protein structures before and after use of our many-body side-chain optimization.

| Model | Optimization | MolProbity<br>Score | Clash<br>Score | Favored<br>Torsions | Rotamer<br>Outliers | Torsion<br>Outliers |
| --- | --- | --- | --- | --- | --- | --- |
| RGN | Minimize | 4.1 | 158.1 | 49.3 | 16.7 | 20.5 |
|  | Sidechains | 3.4 | 74.4 | 52.9 | 9.9 | 9.9 |
| L-RGN | Minimize | 3.3 | 84.5 | 68.3 | 10.2 | 11.8 |
|  | Sidechains | 2.9 | 38.1 | 69.9 | 6.6 | 10.6 |

While both the RGN and Likelihood-RGN testing set structures improve from side-chain repacking, the Likelihood-RGN achieves lower average MolProbity and clash scores than RGN. RGN testing set structures achieve an average MolProbity score of 3.4 and clash score of 74.4; Likelihood-RGN achieves average MolProbity and clash scores of 2.9 and 38.1, respectively. Average MolProbity statistics suggest that the improvement in structure prediction by Likelihood-RGN results in structures that are better suited for biophysical simulations.

## Discussion

In this work, we described an improved dataset and loss function for use in DL approaches to protein structure prediction. We generated the ProteinNetX dataset, which incorporates crystallographic B-factors and computed NMR atomic displacement parameters into the existing ProteinNet protein structure prediction dataset. By reformulating the loss function in the RGN from a least-squares target to a maximum likelihood target, we were able to incorporate these B-factors and atomic displacement parameters as experimental uncertainty measures in model training. Our maximum likelihood model consistently improved network training over a series of trials, both in the initial stability and convergence of model training and in the accuracy of the structures predicted by the final models based on a series of global distance metrics. Likelihood-RGN also predicted structures with more physically realistic backbone torsions, likely a result of the well-defined regions of secondary structure with relatively small B-factors contributing more to model training.

These improvements in secondary structure predictions proved evident in physics-based optimizations. The structures predicted by Likelihood-RGN were more amenable to physics-based optimizations and retained their overall fold better than the structures predicted by the least-squares RGN following energy minimization protocols. This suggests that a maximum likelihood loss model may be better suited for downstream biophysical structural refinement, such as molecular dynamics-based backbone folding(5, 21) or global side-chain optimization(42, 47). Beginning physics-based protein folding from backbones predicted by deep learning, rather than attempting *ab initio* folding, decreases simulation time and improves the resulting structures.

Future directions of our work include improving upon NMR structure coordinate and uncertainty representations in the ProteinNetX, developing a method to compute atomic displacement parameters for single-model NMR and cryo-EM structures, and training a neural network to predict B-factors alongside protein coordinates. Predicted B-factors could help quantify the uncertainty within an individual structure prediction. Knowledge of uncertainty in predicted atomic coordinates would benefit downstream physics-based refinement by guiding optimization methods toward improving lower confidence protein regions.

Our reformulation of RGN’s least-squares loss to a maximum likelihood loss is a novel approach in the effort to apply DL methods to protein structure prediction. The ideas presented here are complementary to existing machine learning approaches to protein backbone folding and can be easily incorporated into other DL models, such as RGN2 and AlphaFold2, to continue to improve our structural coverage of the human proteome. As researchers continue to develop increasingly complex DL models, improvements to loss functions, training datasets, and optimization procedures are imperative to furthering the public effort toward solving the protein folding problem.

## Methods

### A. Curating a Dataset with Atomic Displacement Parameters

#### Existing Protein Datasets for Machine Learning

ProteinNet(24) is a standardized machine learning database of protein sequences, structures, and evolutionary data designed to help develop and assess data driven methods for protein structure prediction. ProteinNet contains six separate datasets that emulate the conditions of prior CASP competitions (CASP7-12). For each CASP, ProteinNet provides a training dataset that contains all protein structures published in the PDB prior to the start date of the competition after filtering out similar or repeated structures based on sequence identity and structures with a poor quality (*e*.*g*., >90% of the sequence being unresolved). The corresponding testing datasets include target sequences and structures from the selected CASP competition. A validation dataset for each CASP was also generated using a clustering process based on sequence identity to mirror the difficulty of the testing dataset. By separating the data in this manner, ProteinNet allows users to directly evaluate their models against results from former CASP competitions and determine algorithmic improvements. The training dataset for our maximum likelihood framework augments the ProteinNet by adding atomic displacement parameters as measures of coordinate uncertainty.

#### Atomic Displacement Parameters

In X-ray crystallography, the B-factor (also called the atomic displacement parameter, Debye–Waller factor, or temperature factor) of an atom describes its vibrational motion about a mean position and thereby influences the X-ray diffraction pattern of the structural model(50). The B-factor is computed using the relationship:

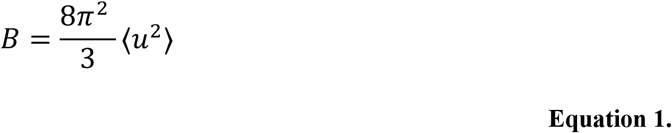

where ⟨*u*^2^⟩ is the mean squared displacement of the atom(51). A large B-factor is correlated with structural regions that have higher flexibility or less certainty in atomic coordinates, whereas a small B-factor is consistent with more rigid, folded regions of a protein structure that have reduced conformational uncertainty. For these reasons, B-factors can serve to indicate uncertainty in protein interatomic distances when training a DL algorithm for structure prediction.

#### Deriving Atomic Displacement Parameters for NMR Structures

Protein structures resolved by NMR spectroscopy lack the defined temperature factors common to structures determined by X-ray crystallography. We derive experimental uncertainties for multi-model NMR structures by computing the root mean square fluctuation (RMSF) of each atom over the NMR ensemble. This per-atom RMSF can be computed in Angstroms *via* the following relationship:

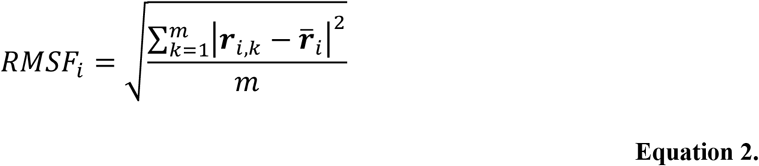

where 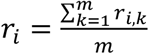 is the average position of an atom over *m* models and *r*_*i,k*_ is the position of atom *i* in the *k*th model(52). Though the dynamics of proteins in solution captured by NMR spectroscopy will differ from the motion of proteins observed in a crystal structure, general trends in uncertainty will remain the same: regions of defined secondary structure will have a lower per-atom RMSF, while this computed value will be much higher in flexible loop regions. To mirror the units and scale of crystallographic B-factors, we multiply each RMSF by a constant factor to obtain the NMR uncertainty value:

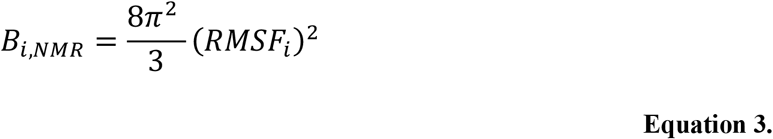

Computing B-factors equivalents for single-model NMR and cryo-EM structures is beyond the scope of this work but can be considered in future work to increase the number of protein structures available in the training dataset.

#### Augmenting the ProteinNet to Include Atomic Displacement Parameters

Using BioJava, a software tool that obtains a protein’s structural information from the RCSB based on its PDB ID(53, 54), we collect B-factors for the backbone atoms of each X-ray crystallography structure in the CASP12 ProteinNet, which is the largest available ProteinNet dataset. We also compute NMR atomic displacement parameters for the backbone atoms of each multi-model NMR structure in the dataset. We compile these structures and atomic displacement parameters, along with the information for each protein included in the original ProteinNet, into a new dataset. We call this augmented dataset the ProteinNetX.

### B. Reformulating Training of the Recurrent Geometric Network Using a Maximum Likelihood Loss Function

The RGN model takes as input an amino acid sequence and its corresponding position specific scoring matrix, and ultimately returns the 3D coordinates of a protein backbone. RGN is comprised of three stages: computation, geometry, and assessment. In the first stage, structural and evolutionary information from amino acids is integrated into adjacent units. Three values are output for each unit, corresponding to the backbone torsional angles (*i*.*e*., φ, ψ, and ω) of each residue. In the second stage, the 3D Cartesian coordinates of the protein backbone are defined by iteratively extending the amino acid chain by one amino acid based on the predicted torsional angles and known bond lengths and bond angles. The third stage outputs the final 3D structure of the protein, evaluates the loss between predicted and experimental structures, and minimizes this loss using backpropagation of the gradient (Figure 1). The loss function used by the original RGN is the distance-based root-mean-square-deviation (dRMSD) metric. The dRMSD is computed by first evaluating the pairwise distances between all atoms in the predicted structure and all atoms in the experimental structure individually, followed by evaluating the RMSD between the two sets of distances.

The ProteinNetX dataset serves as training data for Likelihood-RGN, a modified RGN model that employs the maximum likelihood loss function described below. To derive the loss function, we begin from the electron density *ρ*_*i*_(***d***_*i*_) of an atom at coordinates ***d***_*i*_, which is used to model X-ray diffraction(40)

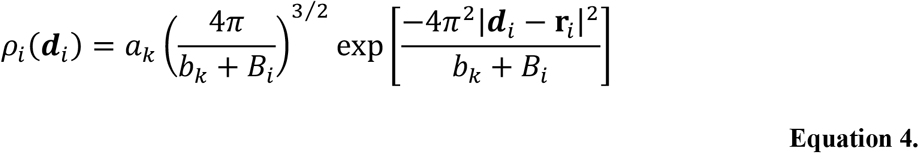

where ***r***_*i*_ are the coordinates of the atomic center, *B*_*i*_ is the atom’s isotropic crystallographic B-factor, and *a*_*k*_ and *b*_*k*_ parameterize a typical Gaussian atomic form factor amplitude and width, respectively(55). Taking the limit as the form factor width goes to zero (*i*.*e*., all electron density is located at the atomic center) and setting the amplitude to unity (*a*_*k*_ = 1), gives the probability of finding the atom at coordinates ***d***_*i*_:

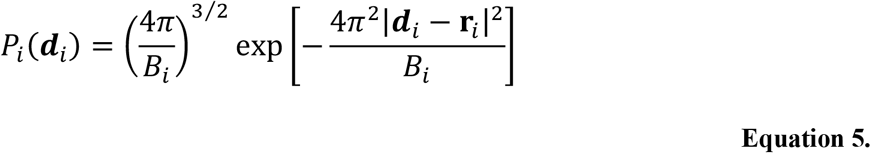

Similarly, a 3D normal distribution with equivalent standard deviations in each dimension {σ = σ_*x*_ = σ_*y*_ = σ_*z*_} and no correlations between dimensions can be modeled by:

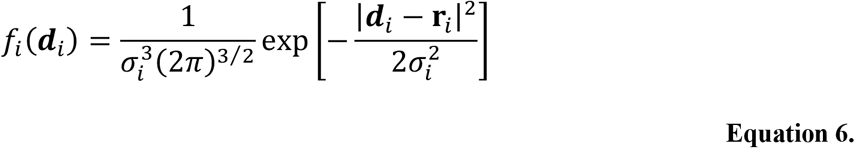

Comparing *f*_*i*_(***d***_*i*_) to *P*_*i*_(***d***_*i*_) shows the variance of Equation 5 is given by

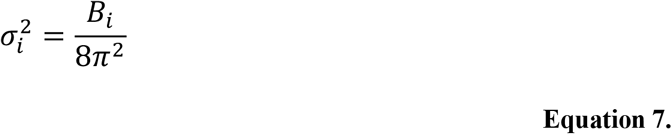

To mirror the dRMSD target used by the original RGN, we now consider a second atom *j* with a measured position ***r***_*j*_ and a variance of 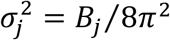. The probability of a predicted atomic separation ***d***_*ij*_ is a Gaussian centered at ***r***_*ij*_ = ***r***_*i*_ – ***r***_*j*_ with a total variance of

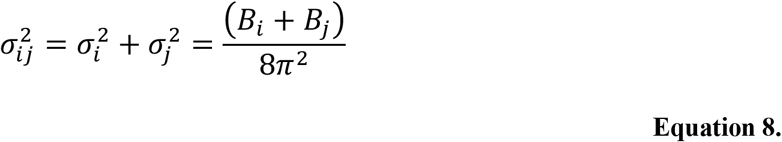

giving the probability density function

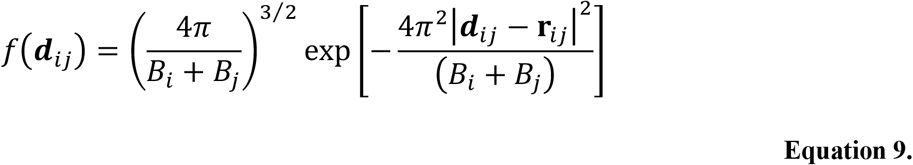

The overall likelihood of the experimental backbone coordinates *X*_*0*_ given the predicted backbone coordinates *X*_*c*_ is then given by a product of interatomic distance likelihoods

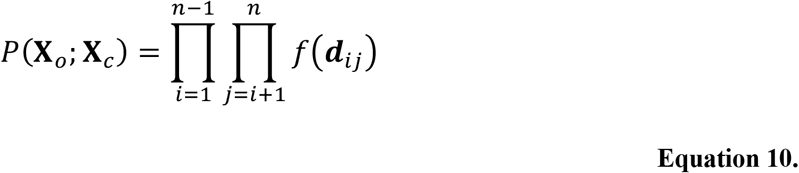

where ***d***_*ij*_ is the predicted atomic separation between the Cα atoms of residue *i* and residue *j* and n is the total number of residues in the protein. The negative of the natural log of the total likelihood then becomes the loss that is minimized during Likelihood-RGN training, given by (ignoring constants):

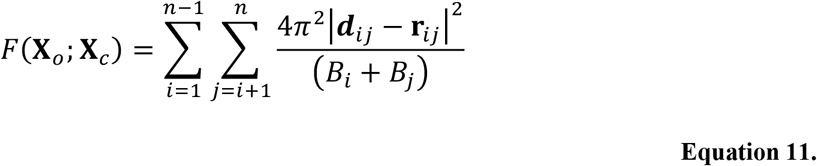

We determine the success of our maximum likelihood loss reformulation by training Likelihood-RGN for protein structure prediction using ProteinNetX and comparing our results to models trained using the original RGN loss function. Hyperparameters for training our model mirror the hyperparameters selected in previous work(29), and the success of model training is initially monitored using the average dRMSD loss of the testing dataset. We then evaluate the accuracy of proteins from the testing dataset predicted by RGN and Likelihood-RGN using a series of geometry metrics, including the RMSD of backbone atoms, GDT(43), GDT-HA(44), TM-Score(45), and favored backbone torsions.

### C. Physics-Based Optimization of Predicted Protein Backbones

We refine the structures of the testing dataset proteins predicted by our two trained models (*i*.*e*., RGN and Likelihood-RGN) with physics-based optimizations using the fixed-charge Amber(12, 13) and the 2018 AMOEBA(46, 56) polarizable force fields with a generalized Kirkwood implicit solvent. We first use the Amber force field to locally optimize the protein structures using the limited-memory Broyden-Fletcher-Goldfarb-Shanno (BFGS) algorithm to a root mean square (RMS) gradient convergence criterion of 0.1 kcal/mol/Å. We then further minimize the structures to an RMS gradient convergence criterion of 0.1 kcal/mol/Å using the AMOEBA force field. Minimizing to a tight convergence criterion using the Amber fixed charge force field first followed by minimization with AMOEBA allows relaxation of the tightly folded predicted backbones, enables a significant reduction in steric clashes, and allows the backbones to find more favorable torsions. This minimization protocol helps prepare the structures for future physics-based simulation and analyses such as molecular dynamics or global side-chain optimization. We evaluate the backbone RMSD, GDT, GDT-HA, TM-Score, and proportion of favored backbone torsions both prior to minimization and after minimization to determine which predicted structures retain their global folds better upon physics-based refinement.

After completing a local optimization, we use our GPU-accelerated many-body optimization(47) to finalize each of the testing set protein structures. Our many-body algorithm builds side chains atoms and computes the global minimum energy conformation for each side chain under the AMOEBA forcefield with generalized Kirkwood implicit solvent. Many-body optimization of side chain atoms dramatically improves the quality of protein structures(47). We evaluate the quality of the testing set structures before and after applying our many-body method using the MolProbity(48, 49) scoring metric, which is widely used by crystallographers to identify high-energy steric clashes and unfavorable side chain or backbone conformations.

## Code and Data Availability

The ProteinNetX datasets, both with and without NMR structures, all code for this work, and the fully trained models are publicly available at https://github.com/SchniedersLab/likelihood-rgn.

## Acknowledgements

All computations were performed on The University of Iowa Argon cluster with support and guidance from Glenn Johnson. Avinash Mudireddy provided guidance on working with TensorFlow and neural networks. GQ was supported by the Barry Goldwater Foundation and the Iowa Center for Research by Undergraduates (ICRU). MRT was supported by the NSF (National Science Foundation) Graduate Research Fellowship under Grant No. 000390183. MJS was supported by NIH R01DK110023, NIH R01DC012049, and NSF CHE-1751688.

## Author Contributions

Developed the theory, G.Q., M.R.T., and M.J.S; performed the experiments, G.Q., M.R.T.; analyzed the data, G.Q., M.R.T., R.A.G., R.J.H.S., M.A., M.J.S.; contributed code, G.Q., M.R.T., M.A., M.J.S.; wrote the manuscript, G.Q., M.R.T., R.A.G., R.J.H.S., M.A., M.J.S.

