## Supplemental Information for "Protein Structure Prediction Using a Maximum Likelihood Formulation of a Recurrent Geometric Network"

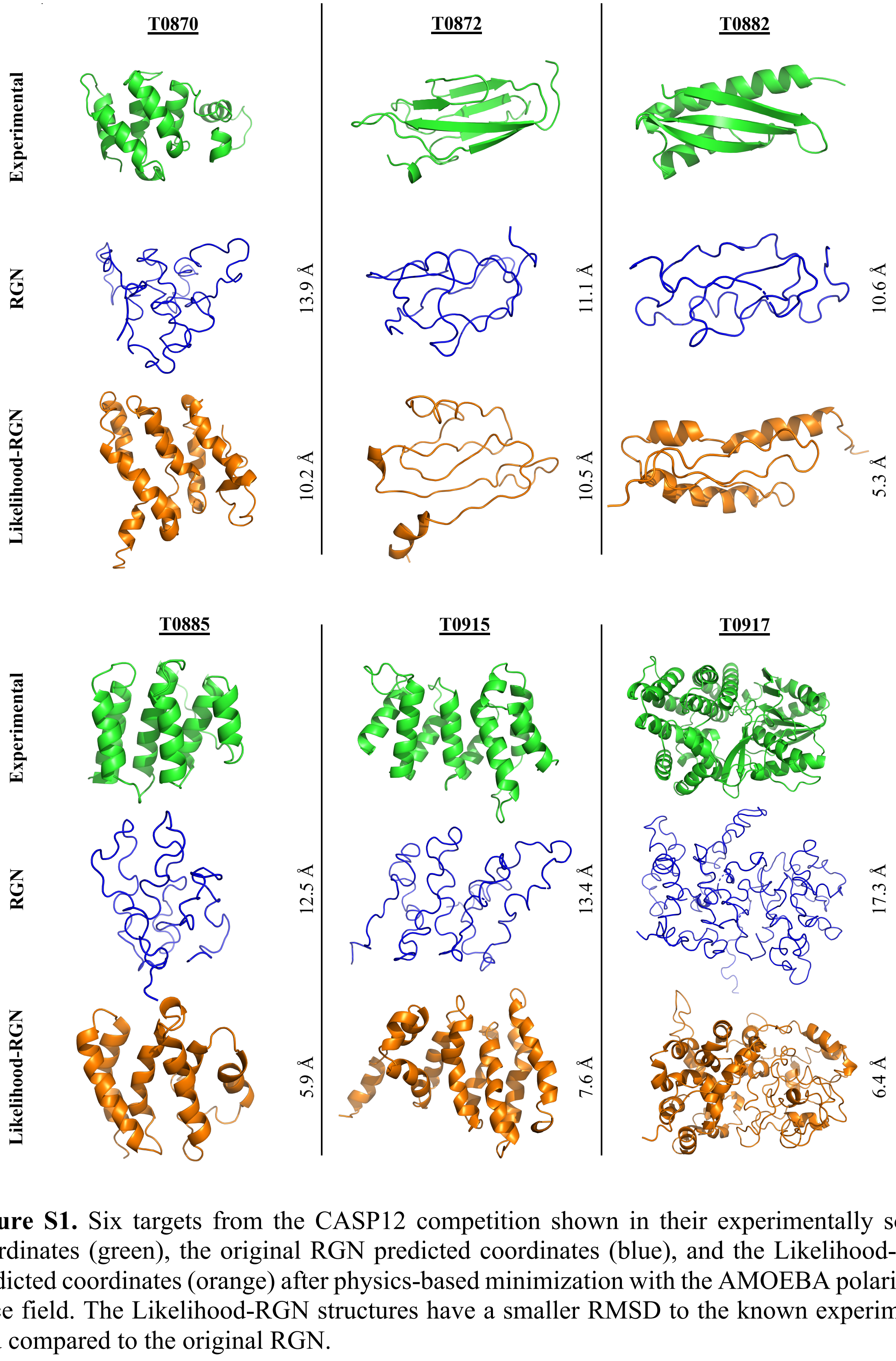


**Figure S1.** Six targets from the CASP12 competition shown in their experimentally solved coordinates (green), the original RGN predicted coordinates (blue), and the Likelihood-RGN predicted coordinates (orange) after physics-based minimization with the AMOEBA polarizable force field. The Likelihood-RGN structures have a smaller RMSD to the known experimental fold compared to the original RGN.

**Table S1.** Average scores across five separate training trials for 63 testing set proteins generated by the RGN (least-squares) and Likelihood-RGN loss functions. The neural networks here were trained on both the X-ray only CASP 12 ProteinNetX dataset and the full CASP 12 ProteinNetX dataset consisting of X-ray and NMR protein structures.

| Training Dataset | Model | dRMSD | RMSD | GDT | GDT-HA | TM-Score | Outlier Torsions | Favored Torsions |
|---|---|---|---|---|---|---|---|---|
| X-ray | RGN | 8.87 | 15.07 | 0.178 | 0.082 | 0.299 | 64.0 | 16.06 |
| | Likelihood-RGN | 8.80 | 14.96 | 0.180 | 0.085 | 0.300 | 50.3 | 23.2 |
| X-ray + NMR | RGN | 8.95 | 15.42 | 0.162 | 0.075 | 0.277 | 58.9 | 21.4 |
| | Likelihood-RGN | 9.01 | 15.04 | 0.169 | 0.079 | 0.286 | 41.4 | 40.3 |

**Table S2.** Average scores over five training trials for the 63 testing set protein structures directly predicted by both RGN and Likelihood-RGN, as well scores for the structures following minimization under the AMOEBA force field. This data was collected from trials that were trained using the X-ray+NMR CASP 12 ProteinNetX dataset.

| Model | Optimization | RMSD | GDT | GDT-HA | TM-Score | Torsion Outliers | Favored Torsions |
|---|---|---|---|---|---|---|---|
| RGN | None | 15.42 | 0.162 | 0.075 | 0.277 | 58.95 | 21.39 |
| | AMOEBA | 16.45 | 0.141 | 0.063 | 0.250 | 27.73 | 45.53 |
| Likelihood-RGN | None | 15.04 | 0.169 | 0.079 | 0.286 | 41.46 | 40.25 |
| | AMOEBA | 16.00 | 0.152 | 0.070 | 0.262 | 22.30 | 53.34 |

**Table S3.** Average scores across five separate training trials for 81 testing set proteins generated by the RGN (least-squares) and Likelihood-RGN loss functions. The neural networks here were trained on both the X-ray only CASP11 ProteinNetX dataset and the full CASP11 ProteinNetX dataset consisting of X-ray and NMR protein structures.

| Training Dataset | Model | dRMSD | RMSD | GDT | GDT-HA | TM-Score | Outlier Torsions | Favored Torsions |
|---|---|---|---|---|---|---|---|---|
| X-ray | RGN | 8.41 | 14.52 | 0.191 | 0.088 | 0.314 | 71.6 | 12.1 |
| | Likelihood-RGN | 8.12 | 14.50 | 0.191 | 0.089 | 0.312 | 66.9 | 17.1 |
| X-ray + NMR | RGN | 8.32 | 14.19 | 0.191 | 0.087 | 0.316 | 48.6 | 26.7 |
| | Likelihood-RGN | 8.17 | 13.63 | 0.206 | 0.097 | 0.335 | 54.1 | 16.3 |
